# IL-10 constrains sphingolipid metabolism via fatty acid desaturation to limit inflammation

**DOI:** 10.1101/2023.05.07.539780

**Authors:** Autumn G. York, Mathias H. Skadow, Rihao Qu, Joonseok Oh, Walter K. Mowel, J. Richard Brewer, Eleanna Kaffe, Kevin J. Williams, Yuval Kluger, Jason M. Crawford, Stephen T. Smale, Steven J. Bensinger, Richard A. Flavell

**Affiliations:** Department of Immunobiology, Yale University, New Haven, CT 06520, USA; Department of Microbiology, Immunology and Molecular Genetics, UCLA, CA 90095, USA; Computational Biology & Bioinformatics Program, Yale University, New Haven, CT, USA; Department of Chemistry, Yale University, New Haven, CT 06520, USA; Institute of Biomolecular Design and Discovery, Yale University, West Haven, CT 06516, USA; Department of Microbial Pathogenesis, Yale University School of Medicine, New Haven, CT 06536, USA; Howard Hughes Medical Institute, Yale University, New Haven, CT 06520, USA

## Abstract

Unchecked chronic inflammation is the underlying cause of many diseases, ranging from inflammatory bowel disease to obesity and neurodegeneration. Given the deleterious nature of unregulated inflammation, it is not surprising that cells have acquired a diverse arsenal of tactics to limit inflammation. IL-10 is a key anti-inflammatory cytokine that can limit immune cell activation and cytokine production in innate immune cell types; however, the exact mechanism by which IL-10 signaling subdues inflammation remains unclear. Here, we find that IL-10 signaling constrains sphingolipid metabolism. Specifically, we find increased saturated very long chain (VLC) ceramides are critical for the heightened inflammatory gene expression that is a hallmark of IL-10-deficient macrophages. Genetic deletion of CerS2, the enzyme responsible for VLC ceramide production, limited exacerbated inflammatory gene expression associated with IL-10 deficiency both *in vitro* and *in vivo*, indicating that “metabolic correction” is able to reduce inflammation in the absence of IL-10. Surprisingly, accumulation of saturated VLC ceramides was regulated by flux through the *de novo* mono-unsaturated fatty acid (MUFA) synthesis pathway, where addition of exogenous MUFAs could limit both saturated VLC ceramide production and inflammatory gene expression in the absence of IL-10 signaling. Together, these studies mechanistically define how IL-10 signaling manipulates fatty acid metabolism as part of its molecular anti-inflammatory strategy and could lead to novel and inexpensive approaches to regulate aberrant inflammation.

## Introduction

IL-10 is an anti-inflammatory cytokine that can limit immune activation in both innate and adaptive immune cell types. IL-10 signaling plays a clear role in modulating mucosal inflammation in the intestine, where deletion of the cytokine, or its receptor (IL-10R), results in inflammatory bowel disease in both mouse and humans^1–5^. Despite IL-10’s importance in maintaining intestinal homeostasis, the exact mechanism of how IL-10/IL-10R signaling reduces inflammation remains unclear. Accumulated work has revealed that inflammatory signals rapidly rewire lipid metabolic programs of immune cells^6–12^ leading us to hypothesize that anti-inflammatory cytokines, such as IL-10, may direct lipid metabolic changes to counteract inflammatory stimuli. Herein, we show that IL-10 regulates sphingolipid metabolism downstream of Toll-like Receptor 2 (TLR2) activation in macrophages. Specifically, in the absence of IL-10 we find increased levels of saturated very long chain (VLC) ceramides which are critical for inflammation both in *vitro* and in *vivo*. Markedly, we find that altered sphingolipid metabolism was mediated by reduced synthesis of mono-unsaturated fatty acids (MUFAs) and could be corrected by providing exogenous MUFAs. Together, these data provide strong evidence that ceramide metabolism is regulated downstream of IL-10R in macrophages by reprogramming de novo synthesis of lipid composition, and helps to explain the molecular circuitry linking alterations in fatty acid homeostasis with inflammation.

### IL-10 signaling regulates sphingolipid metabolism

To better understand global lipid regulation in macrophage, we sought to determine the impact of known anti-inflammatory signaling pathways on TLR-mediated lipidomic shifts^10^. Downstream of many TLRs, macrophages produce IL-10 to limit inflammation in both autocrine and paracrine fashions. To determine if IL-10 signaling can impact macrophage lipid metabolism, we compared lipid profiles of naïve or TLR2 activated WT and IL-10 knockout (KO) bone marrow derived macrophage (BMDMs) using unbiased shotgun lipidomic analysis. Using this strategy, we were able to detect approximately 1,100 lipid species. Principle component analysis indicated that the lipid components of naïve WT and IL-10 KO macrophage were similar, where the quadruplicate samples mostly overlapped for PC1/PC2 (Figure 1A). However, after 48h of TLR2 activation, the lipid composition of WT and IL-10 KO BMDMs diverged (Figure 1A), suggesting that IL-10 signaling is required for normal macrophage lipidomic reprogramming downstream of TLR2 activation. To better understand the differences between WT and IL-10 KO macrophage, we examined what classes of lipids changed in response to TLR2 activation (Figure 1B). Notably, we found significant differences in all measurable subclasses of sphingolipids including ceramides (Cer), modified ceramides (hexosyl (HCer) and lactosyl Ceramides (LCer)) and sphingomyelins (SM), as well as other lipid classes including Triglycerides (TAG), lysophosphatidylcholine (LPCs), and cholesteryl esters (CE). (Figure 1B-D, S1A). Thus, IL-10 is critical for reshaping lipid composition of macrophages in response to pro-inflammatory stimuli.

**Figure 1:**
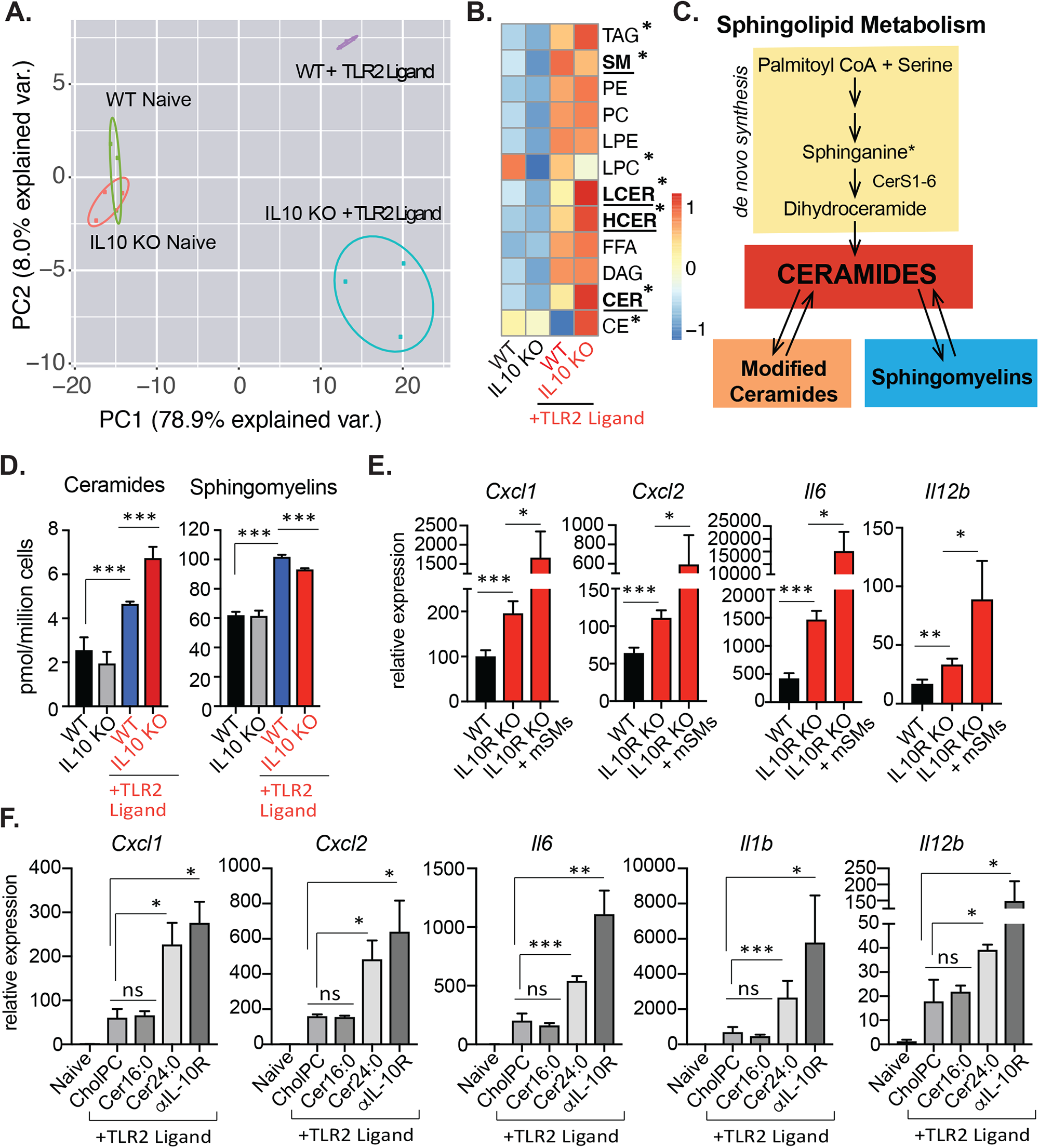
IL-10 signaling regulates sphingolipid metabolism. A) PCA of individual lipids quantified by mass spectrometry (MS) from naïve or TLR2-activated (50ng/mL Pam3CysK4) WT and IL10 KO BMDMs for 48h. Percentage of total variance explained by individual principal components indicated in axis. Prediction ellipses are set at 95% probability (n=4×4 for naïve and 3×3 for TLR2 activated) B) Heat map of individual lipid species measured by direct infusion MS from BMDMs stimulated as in (A). Scaled by row (lipid species). *indicates significant difference (p<0.05) between TLR2 activated WT and IL-10 Kos C) Simplified schematic of sphingolipid metabolism. Ceramides are generated via *de novo* synthesis pathway (yellow box) at the endoplasmic reticulum. Ceramides can be further modified to generate hexosyl and lactosyl ceramides (denoted “modified ceramides”), which can be broken back down into ceramides. Ceramides serve as the building blocks for all sphingomyelin species, which can also be broken down into ceramides. D) Total ceramides and sphingomyelin species measured by direct infusion MS from BMDMs stimulated as in (A) E) qPCR analysis of inflammatory gene expression in WT and IL-10R KO BMDMs activated with TLR2 ligand (50ng/mL Pam3CysK4) for 24h +/-500uM milk-derived sphingomyelins (mSMs) in PBS. (n=3) F) qPCR analysis of inflammatory gene expression in naïve BMDMs or BMDMs activated with TLR2 ligand (50ng/mL Pam3CysK4) for 24h. TLR2-activated macrophage were incubated with Cholesteryl:Phosphatidylcholine (CholPC) lipid sheets alone or with CholPC loaded with ceramide 16:0 (Cer16:0) or ceramide 24:0 (Cer24:0) or cholPC+ anti-IL-10R neutralizing antibody (5ug/mL) for the last 20h of the activation. (n=3 for each group) All experiments are reported as means ± standard deviation (SD) from 3 independent experiments, unless noted otherwise. \**P* < 0.05; \*\**P* < 0.01, \*\*\**P* < 0.005 (two-tailed unpaired Student’s *t* test)

Sphingolipid metabolism is highly complex, where ceramides serve as the central metabolite (reviewed in^13^). Ceramides can be generated via a *de novo* synthesis pathway, and can be further modified to generate complex sphingolipids such as sphingomyelins, and lactosyl/hexosyl-(modified) ceramides. Alternatively, salvage pathways can convert downstream products back to ceramides. (Figure 1C). In line with our previously published data^10^, TLR2 activation strongly induced ceramides, modified ceramides and sphingomyelin species in WT macrophage compared to naïve controls (Figure 1B-D, S1A). Strikingly, we found that loss of IL-10 signaling resulted in increased abundance of total and modified ceramides, while maintaining lower levels of total sphingomyelin (Figure 1B, D, S1A).

Ceramides and sphingomyelins have two fatty acyl tails: the first is 18 carbons long and is derived from the condensation of serine and palmitatoyl-CoA by serine palmitoyltransferase. The second tail is an N-acylated chain of variable length^14^ (Figure S1B). Mammalian cells generate ceramides with second acyl tail lengths of 14-18 carbons (long chain sphingolipids) or 20+ carbons (very long chain sphingolipids) that can be saturated (no double bonds) or mono-unsaturated (one double bond) (Figure S1C). Using our methodology, we were able to consistently quantify sphingolipid species with saturated and mono-unsaturated acyl tails from 14 – 26 carbons. Compared to WT counterparts, all species of ceramides were increased in IL-10 KO macrophage, as well as most modified ceramides (S1D, E). In contrast, most species of sphingomyelins (SMs) were decreased in the IL-10 KO macrophages (Figure S1F). Combined, this data indicates that IL-10 signaling broadly regulates sphingolipid metabolism, largely independent of acyl chain length, via either increased *de novo* synthesis, increased sphingomyelin catabolism or a combination of both.

The impact of sphingolipids on inflammation is controversial. We, and others, speculate that this is due to multiple different pathways capable of reaching the same product and our limited understanding of the mechanisms that regulate sphingolipid *de novo* synthesis versus salvage pathways^15^. IL-10/IL10R deficient macrophage have sustained inflammatory gene expression compared to their WT counterparts in response to TLR2 activation (Figure 1E). Thus, we posited that either the lack of sphingomyelin, the increase in ceramides or both may be important for increased inflammation observed TLR2 activated IL-10/IL-10R KO macrophages. We hypothesized that if lower SM levels found in the IL-10 KO macrophage were important for sustained inflammation, then addition of exogenous SMs would reduce inflammatory gene expression. To test this idea, we generated WT and IL-10R KO BMDMs activated with TLR2 ligands, and added back milk-derived sphingomyelins (mSMs) that consisted of a mixture of long chain and very long SM species. Notably, addition of SMs increased inflammation in the IL-10 KO macrophage (Figure 1E), indicating that reduction of SM in the IL-10 KO was not likely a key component responsible for increased inflammatory gene expression.

Next, we aimed to test if increased ceramides were important for inflammation in IL-10 KO macrophages. As such, we added exogenous ceramides to wildtype macrophages activated with TLR2 ligands. Ceramides, especially long/very long chain ceramides, are extremely hydrophobic, making it difficult to add them to aqueous cell culture mediums. Thus, we generated lipid bilayers comprised of cholesteryl phosphocholine^16^. These bilayers were either left empty (denoted CholPC), or complexed with ceramide 16:0 (Cer16:0, the most abundant long chain ceramide) or ceramide 24:0 (Cer24:0, the most abundant very long chain (VLC) ceramide) (Figure S1D). Addition of Cer16:0 did not induce inflammatory gene expression, whereas addition of Cer24:0, was able to induce inflammatory gene expression to levels similar to those seen in the absence of IL-10/IL-10R signaling (Figure 1F). Combined, these data indicate that loss of IL-10 signaling in macrophages alters sphingolipid metabolism, resulting in increased long chain and VLC ceramides. Addition of exogenous VLC ceramides can induce inflammatory gene expression in WT macrophage to levels similar to IL-10 deficient cells, suggesting dysregulation of VLC ceramide homeostasis may be responsible for exacerbated inflammation that is a hallmark of IL-10 deficiency.

### Genetic inhibition of VLC ceramide synthesis limits inflammation

Ceramides can be synthesized *de novo* by enzymes in the endoplasmic reticulum (ER), or “resynthesized” from degradation products at membranes or in the lysosome (Figure 1C). To understand how IL-10 KO macrophage accumulated ceramides, we performed targeted mass spectrometry (MS) to assess the pool size of sphinganine, a ceramide precursor generated only by the *de novo* synthesis pathway (Figure 1C). Notably, we found that IL-10 deficient macrophage had significantly more sphinganine after TLR2 activation (Figure 2A), suggesting increased *de novo* ceramide synthesis contributed to increased ceramides species found in IL-10-deficient macrophage.

**Figure 2:**
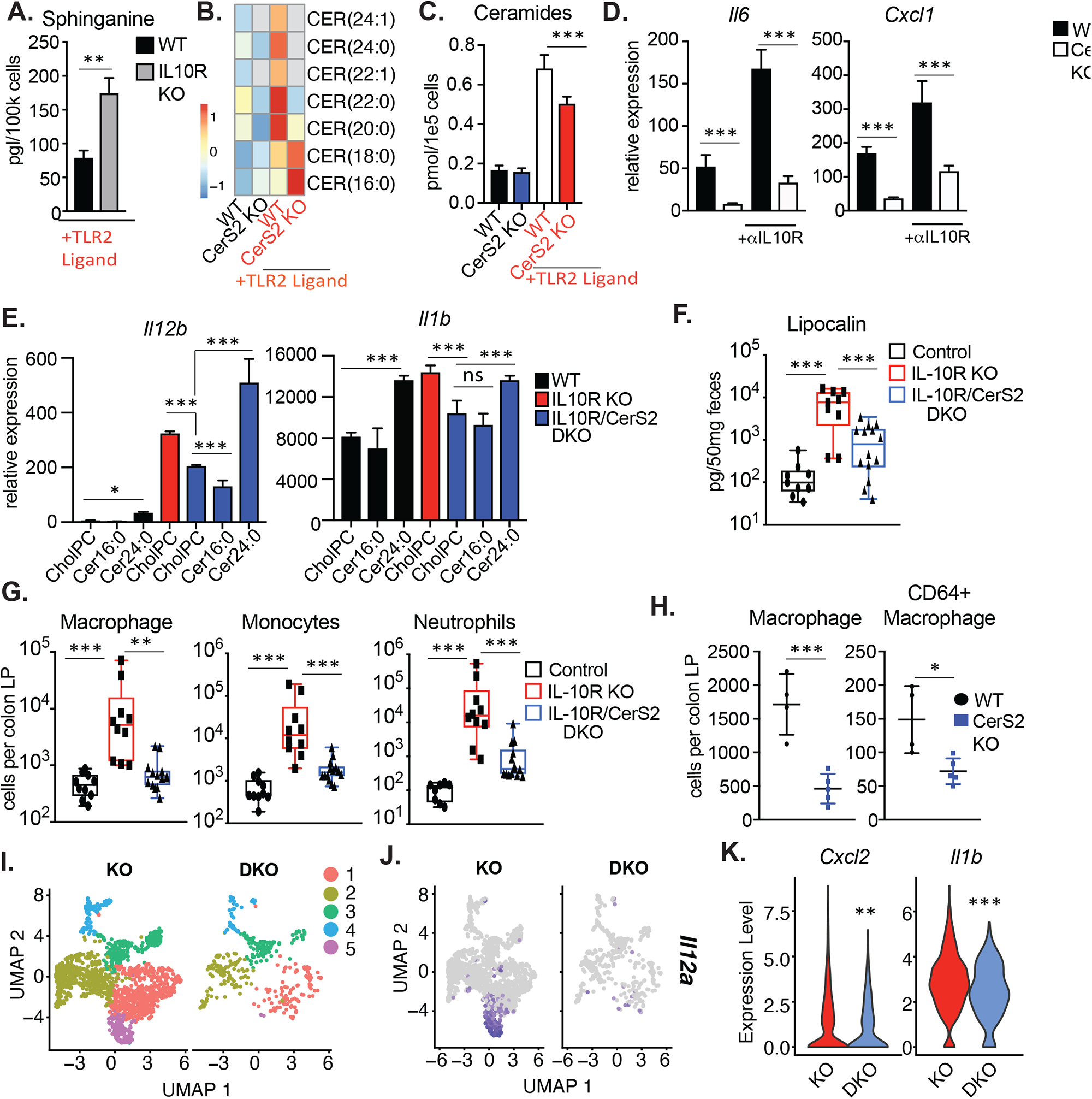
Genetic inhibition of VLC ceramide synthesis limits inflammation: A) MS analysis of Sphinganine in 48h TLR2-activated WT and IL-10R KO BMDMs. (n=3) B) Heat map of individual ceramide species measured by direct infusion MS from naïve or 48h TLR2-activated WT or CerS2 KO BMDMs (n=3 per group) Scaled by row (lipid species) Gray boxes indicated values are zero (below limit of detection). C) Total ceramides species measured by direct infusion MS from BMDMs stimulated as in (B) D) qPCR analysis of *Il6* and *Cxcl1* gene expression in 48h TLR2-activated (50ng/mL Pam3CysK4) WT or CerS2 KO BMDMs +/-anti-IL-10R neutralizing antibody (5ug/mL) for the last 44h of the activation. (n=3 per group) E) qPCR analysis of *Il12b* and *Il1b* gene expression in 48h TLR2-activated (50ng/mL Pam3CysK4) WT (black bars), IL-10R KO (red bar) or IL-10R/CerS2 DKO (blue bars) peritoneal macrophage supplemented with CholPC (vehicle), Cer16:0 or Cer24:0 for the last 44h of activation. (n=3 per group) F) ELISA analysis of lipocalin from feces of control (IL-10R Heterozygous), IL-10R KO and IL-10R/CerS2 DKO chimeric mice (n=9-14) Statistical differences measured by two-sided Mann-Whitney tests G) Macrophage (CD11c+/MHC II+), Monocytes (CD11b+/Ly6C+) and Neutrophils (Cd11b+/Ly6G+) from flow cytometry-based immune cell profiling of the colonic lamina propria (LP) from control (IL-10R Heterozygous), IL-10R KO and IL-10R/CerS2 DKO chimeric mice (n=10-13). Statistical differences measured by Mann-Whitney T-tests H) Macrophage (CD11c/MHCII+) and CD64+ Macrophage from colonic lamina propria (LP) from WT and CerS2 KO chimeric mice (n=4-5) I) scRNAseq UMAP analysis of macrophage clusters from IL-10R KO and IL-10R/CerS2 DKO from cells sorted from the colon lamina propria (LP) J) scRNAseq UMAP analysis of *Il12a* from sorted macrophage from the colon LP of IL-10R KO or IL-10R/CerS2 DKO chimera mice K) scRNAseq gene expression analysis from all clusters of *Cxcl2* and *Il1b* from sorted macrophage from the colon LP of IL-10R KO or IL-10R/CerS2 DKO chimera mice All experiments are reported as means ± standard deviation (SD) from 3 independent experiments, unless noted otherwise. \**P* < 0.05; \*\**P* < 0.01, \*\*\**P* < 0.005 (two-tailed unpaired Student’s *t* test)

The length of the variable acyl tail (Figure S1B) incorporated into ceramides is specified during *de novo* synthesis by a family of enzymes called ceramide synthases (CerS) (Figure 1C). In mouse and humans, there are six ceramide synthases (CerS1-6, formerly called Lass1-6). Utilizing RNAseq data from WT and IL-10 KO macrophage matched to our lipidomics dataset (cells were prepared and stimulated at the same time as cells from the lipidomics experiments in figure 1 but were harvested at 24h post TLR2 activation), we found that BMDMs highly express *Cers2*, *Cers5*, and *Cers6*, but the expression of these enzymes was not influenced by macrophage activation or by IL-10 signaling (Figure S2A). CerS6 regulates the synthesis of long chain ceramides Cer16:0 and Cer18:0, while CerS2 regulates the synthesis of VLC ceramides Cer20:0-Cer26:0. We had observed that Cer24:0 induced inflammation in WT macrophage, thus we generated CerS2 knock-out (KO) mice to determine the impact of VLC sphingolipid synthesis on inflammation. Using lipidomics analysis as in Figure 1, we found that BMDMs from CerS2 KO mice had a variable influence on sphingolipid homeostasis. Specifically, CerS2 KO BMDMs had nearly undetectable levels ceramides with variable acyl chains longer than 20 carbons (Figure 2B, S2B). Despite a significant increase in Cer16:0 (Figure 2B, S2B), which is produced by CerS6, CerS2 KO macrophages had an overall decrease in total cellular ceramides compared to WT controls (Figure 2C). Accordingly, loss of CerS2 dramatically reduced the amount of VLC sphingomyelin species (Figure S2C), however, a striking increase in SM16:0 (Figure S2C) resulted in an overall increase in total sphingomyelins in the CerS2 KOs compared to WT controls (Figure S2D). The pattern of total ceramides and sphingomyelins in the Cers2 KO macrophages is in direct contrast to that seen in the IL-10 KO macrophages, where ceramides are elevated and sphingomyelins are reduced compared to WT controls (Compare Figure 1D to Figure 2C, S2D).

To determine if CerS2 is important for inflammation, we generated macrophage from WT and Cers2 KO mice and activated them with TLR2 ligands alone or in the presence of IL-10R neutralizing antibody. Compared to WT counterparts, CerS2-deficient macrophage showed significantly reduced inflammatory gene expression in response to TLR2 ligands and failed to fully induce inflammatory gene expression during IL-10R blockade (Figure 2D, S2E). This suggests that CerS2 function is important for TLR2-driven macrophage inflammation. To further test this idea, we generated IL10R/CerS2 double KO (DKO) mice. Indeed, loss of CerS2 on the IL-10R deficient background could significantly reduce macrophage inflammatory gene expression (Figure 2E, S2F). Importantly, addition of exogenous Cer24:0, but not Cer16:0, rescued inflammation in the IL10R/CerS2 DKO macrophage (Figure 2E, S2F), indicating that VLC ceramides, the lipid products of CerS2, are important for heightened inflammatory gene expression in IL-10 KO macrophages.

Loss of IL-10 signaling manifests as inflammatory bowel disease in humans and mice, where the majority of inflammatory infiltration and injury is located in the colon^1–3^. Since loss of CerS2 reduced inflammatory gene expression in IL-10R-deficient macrophages, we hypothesized that CerS2 may be important for regulating the colonic inflammation that is the hallmark of IL-10/IL-10R KO animals. We observed that CerS2 KO and CerS2/IL-10R DKO mice were small in stature, had spontaneous seizures, and often died before 10 weeks of age in our colony. This is consistent with phenotypes published by other groups that show that *in vivo* Cers2 deletion results in defects in myelin sheath production and spontaneous seizures^17^. To circumvent complications resulting from these systemic issues, we generated bone marrow chimeras from CD45.2+ IL-10R Heterozygous (Control) mice, IL-10R KO mice and IL-10R/CerS2 DKO littermate mice into CD45.1+ recipients. Microbial variation was minimized by cohousing chimeric mice in a 2x2x2 format for each donor genotype. Bedding was also transferred equally between cages every other week. At 10 weeks post engraftment, fecal samples from the chimeric animals were collected to access lipocalin content, a canonical marker of colonic inflammation^18^. At this timepoint, we observed no overt signs of colonic inflammation (e.g., rectal prolapse or weight loss; data not shown). However, consistent with previous studies in whole body IL-10/IL-10R KO mice^18^, IL-10R KO chimeric mice had significantly higher fecal lipocalin compared to control mice (Figure 2F). Importantly, IL-10R/CerS2 DKO chimeric mice had reduced fecal lipocalin compared to IL-10R KO chimeric mice in both male and female animals (Figure 2F, not shown). To further explore the importance of CerS2 *in vivo*, we collected colon tissue from the chimera mice, isolated the immune cells from the colon lamina propria (LP) and assessed cell populations by flow cytometry (for gating strategies, see Figure S7A, S7B). Consistent with previous studies^3^, when compared to control chimeras, IL-10R KO chimeras had increased inflammatory cells, including macrophage, monocytes, neutrophils, total CD4+ T cells, Th1 CD4+ (IFNψ+) and Th17 CD4+ T (IL-17A+) cells (Figure 2G, S3A). Importantly, loss of CerS2 on the IL-10R-defienct background dramatically reduced immune cell infiltrates compared to the IL-10R KO chimeras (Figure 2G, S3A), indicating that VLC ceramides are important for immune cell mediated colitis driven by the absence of IL-10 signaling.

Colonic inflammation in IL-10-deficient animals is a complicated process that involves crosstalk between several immune cell types and the intestinal epithelium. One hypothesis is that inflammation in the IL-10R KO mice is initiated by increased inflammation from myeloid cells that produce inflammatory cytokines like IL-12, IL-1b, IL-6 and IL-23. These cytokines then polarize T cells towards inflammatory Th1 and Th17 states^19^. In support of this idea, myeloid specific deletion of IL-10R can recapitulate the inflammatory phenotypes found in the whole-body KO, including increased Th1 and Th17 polarization of WT CD4 T cells^20^. To further elucidate how CerS2 impacts inflammation in the colon, we generated WT and CerS2 single KO chimera mice, co-housed the chimeras for 12 weeks and analyzed immune cells from the colonic LP. In this model, there are no drivers of inflammation, allowing us to determine if CerS2 is important for normal immune cell homeostasis in the colon. Notably, we found no reduction in monocytes, neutrophils, total CD4 T cells, Th1 CD4 T cells or Th17 CD4 T cells in the colon LP of CerS2 KO chimeric mice compared to WT controls (Figure S3B). However, CerS2 KO chimeric mice displayed a striking reduction in macrophage (Cd11c/MHCII+) and mature macrophage (Cd11c/MHCII, CD64+) (Figure 2H), suggesting that under basal conditions, CerS2 is important for maintenance of colonic macrophage populations. Combined, our data indicate that CerS2 and VLC ceramides are required for normal macrophage inflammatory activation *in vitro* and *in vivo*.

Colonic macrophages are continually replenished in what’s been dubbed “the macrophage waterfall”^21, 22^ where circulating monocytes migrate to the colon, acquire CD11c, MHCII and eventually CD64 (*Fcgr1*) and become more responsive to TLR signaling, before becoming “tolerogenic” to maintain homeostasis in the colon. As macrophages are turned over, new macrophages mature to repopulate the colon. To determine how loss of CerS2 impacted macrophage homeostasis *in vivo*, we sorted CD45+ immune cells from IL-10R KO or IL-10R/CerS2 DKO chimera animals and analyzed macrophage populations by 10x single cell transcriptomics. UMAP analysis revealed that colonic macrophage isolated from IL-10R KO chimeras grouped into five distinct clusters, where cluster 5 contained highly inflammatory cells (Figure 2I, 2J, S3C, S3D). Strikingly, IL10R/CerS2 DKO chimeras lacked cluster 5, and had reduced expression of inflammatory macrophage markers, cytokines and chemokines compared to IL-10R KO chimeras (Figure 2I, J, K, S3C and S3D). Notably, CerS2-deficient macrophage showed high expression of CD206 (*Mrc1*), suggesting that loss of CerS2 may promote a more tolerogenic macrophage phenotype^23^, compared to IL-10R KO alone (Figure S3C, S3D). Combined, these data indicate that loss of CerS2 on an IL-10R deficient background can mitigate immune cell infiltrates into the colon and reduce *in vivo* macrophage inflammatory gene expression compared to IL-10R KO alone.

### IL-10 induced mono-unsaturated fatty acid synthesis limits ceramide production

To better understand how ceramide biogenesis was regulated in IL-10 KO cells, we examined all known genes required for sphingolipid metabolism from our matched lipidomics RNAseq data set (Figure S2A). And again, we found no difference in gene expression between genotype or activation status (Figure S4A), further supporting the idea that alterations in sphingolipid metabolism driven by IL-10 are not mediated via transcriptional regulation. Interestingly, we found that *Scd2* expression was significantly decreased in IL-10 KO BMDMs (Figure 3A, S3A, bottom row). SCD2 is part of the Stearoyl-CoA Desaturase (SCD) family of enzymes that regulate mono-unsaturated fatty acid (MUFA) synthesis by adding a double bond to saturated fatty acids at the nineth carbon. For example, SCDs convert stearic acid (denoted 18:0) into oleic acid (denoted 18:1). Importantly, we have previously found that SCDs and their products are important in regulating inflammation downstream of TLR activation^10^. To determine if limited *Scd2* gene expression in IL-10 KO macrophage was sufficient to alter *de novo* MUFA synthesis, we employed stable-isotope tracer analysis, where macrophages were fed ^13^C-glucose for 48h with or without TLR2 activation. In line with our previous work, we found that TLR2 stimulation increased *de novo* lipogenesis in WT BMDMs resulting in an increase in total 18:1 and palmitate (16:0) (Figure 3B, C). Strikingly, macrophages lacking IL-10 failed to appropriately reprogram MUFA synthesis downstream of TLR2, resulting in less synthesized 18:1 and less total 18:1 (Figure 3B, C). Palmitate synthesis was unchanged, but total palmitate in the IL10 KO macrophage was higher than WT counterparts (Figure 3B, C), suggesting that altered *de novo* lipogenesis was specific to mono-unsaturated species. To determine if altered MUFA content in IL-10 KOs was important for inflammation, we activated WT and IL-10 KO BMDMs with TLR2 ligand for 48h and added back different free fatty acid species for the last 44h. Importantly, the addition of MUFAs (18:1 and 24:1, an elongation product of 18:1) were able to reduce inflammatory gene expression in the IL-10 KOs, while saturated fatty acids (SFAs; 16:0) and poly-unsaturated fatty acids (PUFAs; 18:2) could not (Figure 3D).

**Figure 3:**
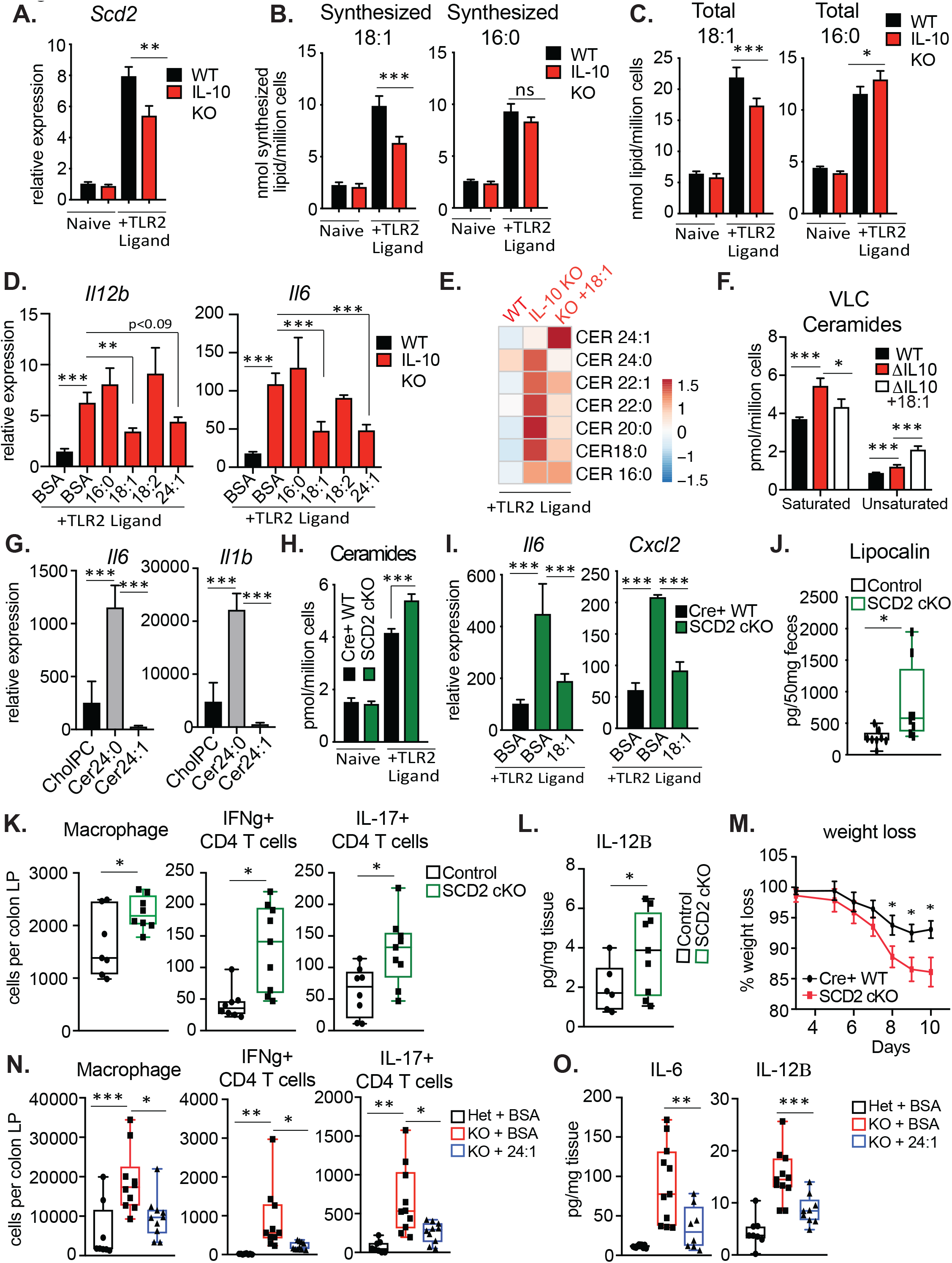
IL-10 induced monounsaturated fatty acid synthesis constrains ceramide production. A) qPCR analysis of *Scd2* gene expression in naïve or 24h TLR2-activated (50ng/mL Pam3CSK4) WT (black bars) or IL-10R KO (red bars) BMDMs (n=3 per group) B) Net synthesis (nmol/million cells) of oleic acid (18:1) and palmitic acid (16:0) as measured by metabolic flux isotope tracer analysis in naïve or 48h TLR2-activated (50ng/mL Pam3CSK4) WT (black bars) or IL-10R KO (red bars) BMDMs (n=4 per group) C) Total cellular (nmol/million cells) of oleic acid (18:1) and palmitic acid (16:0) as measured by GCMS analysis of naïve or 48h TLR2-activated (50ng/mL Pam3CSK4) WT (black bars) or IL-10R KO (red bars) BMDMs (n=4 per group) D) qPCR analysis of *Il12b* and *Il6* gene expression in 48h TLR2-activated (50ng/mL Pam3CSK4) WT (black bars) or IL-10R KO (red bars) BMDMs incubated with vehicle (BSA), or BSA-conjugated 16:0, 18:1, 18:2 or 24:1 for the last 44h of activation. (n=3 per group) E) Heat map of individual ceramide species measured by direct infusion MS from 48h TLR2-activated WT +BSA (vehicle), IL-10 KO + BSA (vehicle) or IL-10 KO + 18:1 BMDMs (n=3 per group). Scaled by row (lipid species) F) Total saturated and unsaturated ceramide species from BMDMs in (E) G) qPCR analysis of *Il6* and *Il1b* gene expression in 48h TLR2-activated (50ng/mL Pam3CysK4) WT BMDMs supplemented with CholPC (vehicle), Cer24:0 or Cer24:1 for the last 44h of activation. (n=3 per group) H) Total ceramides species measured by direct infusion MS from naïve or TLR2-activated Cre+ WT (LysM Cre+/-SCD2^WT/WT^) and SCD2 cKO (LysM Cre+/-Scd2^flox/flox^) BMDMs (n=4 per group) I) qPCR analysis of *Il6* and *Cxcl2* gene expression in 48h TLR2-activated Cre+ WT (LysM Cre+/-SCD2^WT/WT^) BMDMs + BSA, and SCD2 cKO (LysM Cre+/-Scd2^flox/flox^) BMDMs + BSA, and SCD2 cKO BMDMs +BSA-18:1 for the last 44h of activation. (n=3 per group) J) ELISA analysis of lipocalin from feces of Control (LysM Cre+/-) and SCD2 cKO (LysM Cre+/-Scd2^flox/flox^) male mice K) Flow cytometry based immune cell profiling of the colonic lamina propria from naive Control (LysM Cre+/-) and SCD2 cKO (LysM Cre+/-Scd2^flox/flox^) male mice L) ELISA analysis of IL-12B (IL12p40) from colonic explants of naïve Control (LysM Cre+/-) and SCD2 cKO (LysM Cre+/-Scd2^flox/flox^) male mice incubated ex vivo for 24h. M) Weight loss measured during DSS-induced colitis in Control (LysM Cre+/-) and SCD2 cKO (LysM Cre+/-Scd2 flox/flox) female mice (n=9 per group) error bars represent Standard Error of the Mean (SEM). N) Flow cytometry based immune cell profiling of the colonic lamina propria from IL-10R heterozygous (het) mice gavaged with BSA, IL-10R KO mice gavaged with BSA or IL-10R KO mice gavaged with BSA-24:1 (all male mice) (n=9-10) O) ELISA analysis of IL-6 and IL-12B (IL12p40) from colonic explants from IL-10R heterozygous (het) male mice gavaged with BSA, IL-10R KO male mice gavaged with BSA or IL-10R KO male mice gavaged with BSA-24:1 incubated ex vivo for 24h. (n=10-12) All experiments are reported as means ± standard deviation (SD) from 3 independent experiments, unless noted otherwise. \**P* < 0.05; \*\**P* < 0.01, \*\*\**P* < 0.005 (two-tailed unpaired Student’s *t* test)

Next, we sought to determine if ceramide biogenesis is influenced by *de novo* MUFA synthesis. To test if altered MUFA content in IL-10 KO macrophage could alter ceramide species, we used shotgun lipidomic analysis to examine how ceramide species changed IL-10 KO BMDMs in response to oleic acid (18:1) addition. While addition of 18:1 did not lower total ceramide abundance in the IL-10 KO macrophages (Figure S4B), it significantly lowered all saturated VLC ceramides, including Cer24:0 (Figure 3E, F). Notably, addition of 18:1 had no effect on Cer16:0 and increased the amount of Cer24:1 and total mono-unsaturated ceramides (Figure 3E, F). This led us to ask if saturated and unsaturated VLC ceramides influenced inflammation in the same way. Indeed, unlike Cer24:0, Cer24:1 did not induce inflammatory gene expression in macrophage (Figure 3G), indicating that only saturated VLC ceramides can drive inflammation.

In our previous studies, we found that inhibition of MUFA synthesis prolonged inflammatory gene expression in macrophage^10^, but the mechanism of action behind this phenotype was unclear. To determine if MUFA synthesis was important for macrophage ceramide content, we acutely inhibited SCD activity with a small molecule inhibitor (denoted SCDi) in combination with TLR2 activation. Importantly, we found that SCDi treatment increased total ceramides and saturated ceramides, while not altering the pool size of unsaturated ceramides (Figure S4C in the IL-10 KO, the observed increase in saturated VLC ceramides and inflammatory gene expression produced by SCDi could be mitigated by the addition of 18:1 (Figure S4C, S4D).

Genetic deletion of *Scd2* is embryonic lethal in mice, thus, to better understand the impact of SCD2 on inflammation *in vivo*, we generated *Scd2*^flox/flox^ mice and crossed them to the LysM-Cre+ mice to generate animals with myeloid-specific deletion of SCD2 (denoted SCD2 cKO). Importantly, we found that genetic ablation of *Scd2* phenocopied IL-10 deficiency, where SCD2 cKO macrophage had increased ceramides (Figure 3H) and increased inflammatory gene expression in response to TLR2 activation that could be reduced by the addition of 18:1 (Figure 3I). In line with this *in vitro* data, SCD2 cKO mice had basal elevation of fecal lipocalin compared to cohoused Cre+ littermate controls (Figure 3J). Additionally, SCD2 cKO mice had increased macrophages in the colon without impacting neutrophil or monocyte populations (Figure 3K, S4E). Strikingly, SCD2 cKOs also displayed increased total CD4+ T cells, IFNψ+ and IL-17A+ CD4+ T cells (Figure 3K, S4E), indicating that myeloid-specific SCD2 expression could impact T cell activation in the colon. In line with this idea, we found that *ex vivo* colonic explants from SCD2 cKO mice produced more soluble IL-12B (IL-12p40) compared to tissue collected from cohoused Cre+ littermate controls (Figure 3L). We hypothesized that the increased basal inflammation found in the SCD2 cKO mice might predispose these animals to models of chemically-induced colitis, such as Dextran Sodium Sulfate (DSS). Indeed, after 5 days of DSS treatment, followed by 7 days of recovery, we found that SCD2 cKO mice lost significantly more weight than the cohoused littermate Cre+ control mice, and had significantly increased numbers of infiltrating myeloid and lymphoid immune cells in the colon LP (Figure 3M, S4F, S4G). In sum, the phenotypes observed in SCD2-deficient macrophages and mice recapitulate phenotypes in macrophages and mice lacking IL-10 signaling. Together, these findings suggest that perturbed MUFA synthesis is a major underlying cause for accumulation of saturated ceramides and heightened inflammation in IL-10 deficiency.

Since exogenous MUFAs could limit inflammation in IL-10 deficient macrophage, and loss of myeloid-specific SCD2 could induce colonic inflammation, we asked if exogenous MUFAs could reduce colonic inflammation in the IL-10/IL-10R KO mice. To test this, we started with two groups of IL-10R KO mice and two groups of IL-10 KO mice that had similar levels of colonic inflammation, measured by fecal lipocalin content (Figure S5A, S5B). For each genotype, one group of mice was orally gavaged with BSA, the carrier protein used to solubilize free fatty acids, or with BSA conjugated to either 24:1 (IL-10R KOs) or 18:1 (IL-10 KOs). Mice were gavaged for 14 days before colonic tissue was collected for immune cell analysis and cytokine secretion. Strikingly, we found that 24:1 or 18:1 treatment significantly reduced fecal lipocalin compared to BSA alone (Figure S5A, S5B). Furthermore, MUFA gavage reduced myeloid and lymphoid immune cell populations in the colon and reduced IL-6 and IL-12B secretion from colonic explants (Figure 3N, O, S5C, S5D), at levels similar to ablation of CerS2 (Figure 2G, H). Combined, these data further suggest that MUFA synthesis serves as a negative feedback mechanism downstream of IL-10 signaling to dampen inflammation, and that MUFA synthesis is important to control intestinal inflammation.

## Discussion

In this study, we mechanistically delineate crosstalk between innate immune regulation, lipid metabolism and intestinal homeostasis. Here, we uncover a previously undescribed mechanism by which the anti-inflammatory cytokine IL-10 regulates the duration of inflammation. Specifically, we find that IL-10 signaling regulates sphingolipid homeostasis by limiting *de novo* ceramide synthesis to modulate inflammatory gene expression. Unexpectedly, we found that altered sphingolipid production was not mediated at the level of transcriptional regulation of the pathway enzymes (Figure S4A). Instead, changes in ceramide biogenesis were traced to IL-10’s ability to induce synthesis of long chain MUFAs. These data suggest that the *de novo* ceramide synthesis pathway can be mediated by the enzymatic activity of SCD2 (Figure 3H), and that ceramide production can be influenced by IL10-driven fatty acid desaturation. To date, there are few known regulatory checkpoints for *de novo* ceramide synthesis, and how limited MUFA synthesis regulates this pathway remains unclear. One attractive possibility is that when MUFA synthesis is limited, this results in 1) alterations in the pool size of available fatty acid-CoA, like 24:0-CoA and 24:1-CoA, that are utilized to generate VLC mono-unsaturated ceramides and 2) an increase in saturated fatty acid precursors such as palmitate. The condensation of palmitate and serine if the first step in ceramide *de novo* synthesis pathway and is required for the generation of all downstream sphingolipid metabolites. Indeed, we find IL-10 deficiency increases total cellular palmitate in macrophage (Figure 3C), but further biochemical analysis will be required to fully delineate how MUFAs impact ceramide *de novo* synthesis.

Markedly, these are the first observations that exogenous addition of VLC, but not long chain, ceramides can regulate macrophage inflammation (Figure 1F), although other groups had suggested these outcomes *in silico*^24^. Significantly, we find that the VLC ceramide tail “desaturation status” is important for induction of inflammation (Figure 3G), and future studies will focus on how saturated but not unsaturated VLC ceramides mediate inflammatory gene expression. Additionally, we find that loss of IL-10 signaling reduces most saturated and all mono-unsaturated SMs. This could either be due to perturbed ER-to-golgi transport that is required for SM synthesis or due to enhanced SM degradation. The latter option has been observed in response to inflammatory cytokine signaling in many immune cell types, but the exact mechanism for how increased SM degradation impacts inflammation remains controversial. As such, we find that a mixture of long chain and VLC SMs can enhance inflammatory gene expression in macrophage, but it is unclear if this enhanced inflammation is mediated via the same pathways we find responsible for VLC ceramide-induced inflammation.

Finally, we find that genetic reduction of VLC ceramides or oral gavage with MUFAs can reduce colonic inflammation found in IL-10/IL10R deficient mice (Figure 2F-L, Figure 3N, O), suggesting that *de novo* lipid synthesis pathways are important to control intestinal inflammation. Humans with deleterious mutations in IL-10, or its receptor, present with severe, life-threatening enterocolitis within the first few months of their lives. These patients fail to respond positively to standard forms of immune-suppression, such as anti-TNF biologicals or steroids. Outside of hematopoietic stem cell transplant, there is no effective treatment option for inflammatory bowel syndrome in humans who lack functional IL-10 signaling. Intriguingly, dietary intervention with the Mediterranean Diet, which is high in unsaturated fats, has been shown to reduced disease parameters in patients with IBD in as little as 6 months^25^. A key component of the Mediterranean Diet is olive oil, which is high in MUFA content. Thus is it tempting to speculate that the benefits of the Mediterranean Diet may dependent in part on MUFA-mediated repression of colonic inflammation. Indeed, our data strongly suggests that oral delivery of exogenous MUFAs is beneficial to curb inflammation in the colon and may serve to establish new treatments for colitis focused on fatty acid “metabolite correction”.

## Acknowledgements

We would like to thank Jon Alderman, Cynthia Hughes, Lin
da Evangelisti, Elizabeth Hughes-Picard, and Beth Cadugan for their help with technical and administrative duties.

## Funding

A.G.Y. is supported by the Howard Hughes Medical Institute Hanna H. Gray Fellowship. W.K.M. is supported by NIH T32-DK007356, and is the recipient of a Cancer Research Institute/Irvington Postdoctoral Fellowship.

## Author Contributions

A.G.Y. conceptualized of this project, designed/implemented experiments, analyzed data, provided funding, and constructed the manuscript. MHS designed and implemented experiments, and analyzed data. QZ KJW, WH, WM, RB, and JO designed and implemented experiments. RQ provided formal lipidomics analysis, RNAseq analysis and 10x scRNAseq analysis. UK, JC, STS, SJB and RAF provided resources/supervision/funding for this project, contributed to conceptualization and revision of the manuscript.

## Competing Interests

None. But we would like to disclose that RAF is an advisor to Glaxo Smith Kline.

## Data Availability

The datasets generated during the current study are available from the corresponding author on reasonable request. Upon Publication, all datasets will be made publicly available on NCBI GEO.

## Materials and Methods

### Mouse Strains

IL-10 KO (JAX #002251) and IL-10R KO (JAX #005027) mice were purchased from Jackson labs. Both KO strains were crossed to WT *C57BL/6* (JAX 000664) to generate heterozygous mice. All future cohorts of mice were generated from het x het breading to minimize microbial diversity. Cohorts of mice were aged in cohoused cages with 3x WT and 3x KO unless noted otherwise. LysM-Cre^+/-^ (JAX 004781) mice were also purchased from Jackson Laboratory.

SCD2 flox/flox mice and CerS2 KO mice were generated in our in-house CRISPR Core and are available upon request. For Bone marrow chimeras, donor bone marrow was transferred into CD45.1 PepBoy (Jax #002014) recipient mice. All animal experimentation was performed in compliance with Yale Institutional Animal Care and Use Committee protocols.

### Mouse cells

Bone marrow was differentiated into macrophages in DMEM containing 10% FBS (Sigma), 5% M-CSF conditioned media or 50ng/mL M-CSF (Biolegend) (results did not differ), 1% pen/strep (Gibco), 1% glutamine (Invitrogen) 0.5% sodium pyruvate (Invitrogen) for 7-9 days prior to experimental use. Cells were changed to media with 5% FBS at the time of TLR stimulation. Peritoneal macrophages were harvested after 96h thioglycallate-treatment. Macrophages were washed, counted, plated and immediately activated by TLR agonists.

### Reagents

Cells were treated with 50ng/mL Pam3CSK4 (Invivogen tlrl-pms). CholPC (#700123), Cer16:0 (#860516), Cer24:0 (#860524), Cer24:1 (#860525), and milk derived SMs (#860063P) were purchased from Avanti Lipids.

#### Preparation of Ceramide:Cholesteryl Phosphocholine (CholPC)

Ceramides and CholPC were dissolved into 100% chloroform to 12.5mM. 1:1 mixtures of ceramide and CholPC were mixed and dried under nitrogen. Lipids were rehydrated in PBS to 3.125mM Cer:3.125mM CholPC (or CholPC:blank), and sonicated at 55 degree C for ∼1h until all precipitates were in solution. Lipid mixtures were added to cell cultures at 1:100 dilution for 31.25uM final concentration for each ceramide species.

#### Preparation of 24% BSA

FFA-free, endotoxin-low BSA (Sigma A8806) was gradually add 6g of fatty-acid-free, endotoxin-low BSA (Sigma A8806) to 17.5mL 150mM NaCl in a beaker at 37°C while stirring. This takes about 6-7h. Add one “scoop” of BSA using a small spatula. Wait 3-5 mins until BSA is completely dissolved before adding next scoop.) 24% BSA pH was adjusted to 7.4 with 1N NaOH. Add 1M NaOH slowly in 20uL increments, while stirring. Repeat 10x times. Wait 1-2mins and pH. Repeat in 10x increments slowly until desired pH is achieved. Do not add higher volume or use more concentrated NaOH. This will cause BSA to crash out of solution.

#### Preparation of BSA-conjugated Free Fatty Acids

Free fatty acids (FFAs: Nucheck Prep) were dissolved 150mM NaCL pH=7.4 to 25mM final concentration (ie 50mg 18:1 was dissolved into 7mLs 150mM NaCl + 48uL NaOH). Mixture was shaken vigorously, and heated at 65 deg C until in solution). 24% BSA solution (ice cold) is finally combined with 25mM oleic acid solution (room temp) in a 54:46 ratio to yield approximated 12.5mM final concentration with pH=7.4. Mix for 10 mins at RT on shaker before use. Store at -20 deg C.

#### Addition of FFAs into cell culture

FFAs were diluted into cell culture medium at 1:500 dilution for 25uM final concentration. FFA’s were added 4h post TLR stimulation.

#### Oral gavage of BSA-18:1

Mice (15-18g, 5-6weeks old) were gavaged for 14 days with 100uL of BSA alone or 6.25uM BSA-18:1 (preparation described above, diluted 1:1 with PBS)

#### Inhibition of MUFA synthesis by SCDi

SCDi (Cay10566; Fisher NC0493687) was dissolved into DMSO to a final concentration of 10uM. This solution was used immediately, only once, and did not undergo any freeze/thaw cycles (any freeze/thaw cycles killed the activity of this compound as measured by isotope tracer analysis). Working solution was generated by 1:1000 dilution for a final working solution of 10nM.

#### ELISAs

Lipocalin (R&D), IL-6 (R&D) and IL12B (Biolegend) ELISAs were preformed to manufacturers instructions

### Lipidomics Analysis

Macrophages were cultured in 6 well dishes (Fisher 08-772-1B) and stimulated with TLR ligands as described above. 48 hours post-stimulation, cells were imaged for cell count as previously described (York, 2015), scraped and spun down in PBS, and snap-frozen as cell pellets. A modified Bligh and Dyer extraction (Bligh, 1959) was carried out on samples. Prior to biphasic extraction, a 13-lipid class Lipidyzer Internal Standard Mix is added to each sample (AB Sciex, 5040156). Following two successive extractions, pooled organic layers were dried down in a Genevac EZ-2 Elite. Lipid samples were resuspended in 1:1 methanol/dichloromethane with 10mM Ammonium Acetate and transferred to robovials (Thermo 10800107) for analysis. Samples were analyzed on the Sciex Lipidyzer Platform for targeted quantitative measurement of 1100 lipid species across 13 classes. Differential Mobility Device on Lipidyzer was tuned with SelexION tuning kit (Sciex 5040141). Instrument settings, tuning settings, and MRM list available upon request. Data analysis performed on Lipidyzer software. Quantitative values were normalized to cell counts. PCA and heatmaps were generated using guidelines described in ^26^

### Detection of Sphinganine

The detection and quantification of sphinganine (SPA) were performed using an Agilent 6490 ESI-QQQ-MS/MS instrument fitted with an electrospray ionization (ESI) source (positive) coupled to an Agilent 1290 Infinity HPLC system. The target lipid was analyzed utilizing a gradient program on a Phenomenex Kinetex 1.7 mm C18 100 Å (100 X 2.1 mm) (10% → 100% MeCN in water with 0.1% formic acid for 7 min, 0.3 mL/min, column temperature: 50°C). The detection was conducted with the optimized collision energy at 20V and facilitated by dynamic multiple reaction monitoring (MRM) mode with the mass transition *m/z* 302→ 284.

### Isotope Enrichment Experiments

Day 8 differentiated BMDMs were transferred to complete media containing 50% U^13^C-glucose with or without TLR stimulation for 48hr before collection. See ^9^ for further details. Analysis of labeled fatty acids and cholesterol was performed as described in ^27^. The relative contributions of synthesis to the total cholesterol pool over the 48h-labeling period were determined by fitting the isotopologue distributions for cholesterol in a model similar to Isotopomer Spectral Analysis (ISA) as described in ^27^

### Gene expression analysis

RNA was extracted from all cells with Trizol using manufacturer’s protocols. cDNA was synthesized with Applied Biosystems High Capacity cDNA Synthesis Kit as per manufacturer’s instructions (700ng/uL RNA per cDNA synthesis reaction). Quantitative PCR (qPCR) was conducted on the BioRad qPCR machine using SYBR Green Master Mix (BioRad) and 0.5 μmol/L primers. Relative expression values are normalized to control gene (rRNA 36B4) and expressed in terms of linear relative mRNA values. Primer sequences are available upon request.

### Fecal Lipocalin preparation

Feces of each mouse was collected, weighed and dissolved in PBS+0.1% Tween. Fecal pellets were disrupted using a tube shaker for 5 mins at fully speed. Samples were spun down and the resulting supernatants were diluted 1:10 and utilized for lipocalin ELISAs.

### DSS Colitis

Mice were given 1.5% DSS in their drinking water for 5 days, before changing back to normal water bottles. Mice were weighted through the duration of the experiment and sacrificed at Day 12.

### Isolation of immune cells from the colon LP

Colons were collected, flushed with 10mL PBS, and split length wise, and rinsed again in PBS. The epithelia fraction was removed with two 10mL washes of Epi Wash Buffer (1x HBSS, 5mM EDTA, 1mM DTT) at 37 deg C for 20 mins while shaking at 220RPMs. Colons were removed from Epi Wash Buffer, washed in PBS and finely minced with a razor blade. Minced tissue was transfer into 5mL Digestion buffer (1x DMEM, 5% FBS 1mg/mL Collagenase D, 0.5mg/mL DNAase) and incubated at 37 deg C for 60 mins while shaking at 220 RPMs. After digestion, samples were strained though at 70um filter and washed 2x 20mL with 1x DMEM + 5% FBS. Cells were then divided into 5 groups for different staining panels. Cells were stained with antibodies and fixable viability dye at 4 deg C for 30 mins in FACS buffer (2% FBS in PBS), washed 2x, and run on an LSR2 or fixed (BD Cytofix/CytoPerm #554722) for intracellular staining. AccuCheck cell counting beads (Invitrogen PCB100) were utilized for cell number quantification

### Flow Cytometry antibodies

From Biolegend: CD45.2 (Clone SJL), CD45.1 (clone A20), CD11b (Clone M1/70), CD11c (Clone N418), MHC II (I-A/I-E, clone M5/114.15.2),CD64 (clone X54-5/7.1), Epcam (clone G8.8), Ly6G (clone 1A8), Ly6C (clone HK1.4), TCRb (clone H57-597 ), CD4 (clone RM4-5), CD8a (clone 53-6.7), IL-17A (clone TC11-18H10.1), IFNg (clone XMG1.2), Fixable Viability Dye BV510 (BD #564406). For gating, see Figure S7A&B.

### 10x scRNAseq sample preparation

Isolated cells from the colons were sorted for CD45+ CD11b/CD11c/CD64+ cells and TCRB+ cells to exclude neutrophils and B cells. Cell populations were combined with 5,000 of each population for 10,000 cells final. Cells were further prepared using the 10x Single Cell comptroller per manufacturer’s instructions.

### scRNAseq data analysis

Cellranger v6.0.1 was used to align scRNA-seq samples to the mouse genome (mm10). Data analysis was performed using Seurat v4.3.0 R package^28^, including cell-type identification and comparative analyses between conditions. In the step of quality control, poor-quality cells with the number of expressed genes < 300 or > 5000 were filtered out. We also excluded cells if their mitochondrial gene percentages were over 10%. After combining cells from all samples, we first normalized the raw count matrix and then defined 2000 top variable genes. We then applied principal component analysis (PCA) for dimensionality reduction and retained 30 leading principal components for cell clustering. Specifically, the Shared Nearest Neighbor (SNN) graph was constructed by calculating the Jaccard index between each cell and its 20-nearest neighbors, which was then used for cell clustering based on Louvain algorithm (with a resolution of 0.3). After identifying cluster-specific genes, we annotated cell types based on canonical marker genes (Lyz2 for macrophage) and focused on macrophage populations in the downstream analyses. To test the differential expression of inflammatory genes between IL-10R KO and Cers2/IL-10R DKOs, we first applied ALRA^29^ to impute the gene expression profiles and then used the non-parametric Wilcoxon rank sum test to get their p-values.

### RNAseq data analysis

Adapter sequences were removed from the raw sequencing reads using the tool Cutadapt (https://journal.embnet.org/index.php/embnetjournal/article/view/200). STAR^30^ v2.5.3 was then used to align the trimmed sequencing reads to the mouse genome (mm10) with default parameters. After sorting the generated SAM files (as the output of alignment) with Picard Toolkit (Picard Toolkit.” 2019. Broad Institute, GitHub Repository. https://broadinstitute.github.io/picard/; Broad Institute), we counted the number of reads mapped to each gene using HTSeq^31^ v0.6.1. Subsequently, we employed DESeq2^32^ v1.26.0 R package to identify significantly differential genes between naïve and TLR2-activated (24h) WT and IL-10 KO BMDMs.

## SUPPLEMENTAL FIGURE LEGENDS

**Supplemental Figure 1:**

A) Total Cholesteryl Esters (CE), Diacylglycerols (DAG), Free Fatty Acids (FFAs),Hexosyl Ceramides (HCer), Lactosyl Ceramides (LCer), Lysophosphatidylcholine (LPCs), Lysophosphatidylethanolamine (LPE), phosphatidylcholine (PC), phosphatidylethanolamine (PE) and triglycerides (TAGs) measured by direct infusion MS from naïve or 48h TLR2 activated WT and IL10 KO BMDMs

B) Schematic of Ceramide 16:0 and Sphingomyelin (SM) 16:0. Both sphingolipids contain an 18 carbon long constant chain (top, black numbers), and a 16 carbon long variable chain (bottom, red numbers)

C) Schematic of saturated (Cer24:0) versus unsaturated (Cer24:1) ceramides. Double bond in 24:1 is located at the arrow.

D) Ceramide species measured by direct infusion MS from WT and IL10 KO BMDMs that are naïve or activated with TLR2 ligand (50ng/mL Pam3CysK4) for 48h.

E) Heat map of individual hexosyl ceramide (HCer) species measured by direct infusion MS from WT and IL10 KO BMDMs that are naïve or activated with TLR2 ligand (50ng/mL Pam3CysK4) for 48h. Scaled by row (lipid species) *indicate significance of p<0.05 between WT and IL-10 KO activated with TLR2

F) Heat map of individual sphingomyelin (SM) species measured by direct infusion MS from WT and IL10 KO BMDMs that are naïve or activated with TLR2 ligand (50ng/mL Pam3CysK4) for 48h. Scaled by row (lipid species) *indicate significance of p<0.05 between WT and IL-10 KO activated with TLR2

All experiments are reported as means ± SD from 3 independent experiments, unless noted otherwise. \**P* < 0.05; \*\**P* < 0.01, \*\*\**P* < 0.005 (two-tailed unpaired Student’s *t* test)

**Supplemental Figure 2:**

A) RNAseq analysis of Ceramide synthase genes naïve or TLR2 activated WT and IL10 KO BMDMs from RNAseq analysis from samples matched to lipidomics analysis in Figure 1 (TPMs must be above 0 to be listed). (n=2)

B) Ceramide species measured by direct infusion MS from naïve or 48h TLR2-activated WT or CerS2 KO BMDMs

C) Sphingomyelin (SM) species measured by direct infusion MS from naïve or 48h TLR2-activated WT or CerS2 KO BMDMs

D) Total Sphingomyelin (SM) measured by direct infusion MS from naïve or 48h TLR2-activated WT or CerS2 KO BMDMs

E) qPCR analysis of *Il1b* gene expression in 48h TLR2-activated (50ng/mL Pam3CysK4) WT or CerS2 KO BMDMs +/-anti-IL-10R neutralizing antibody (5ug/mL) for the last 44h of the activation. (n=3 per group)

F) qPCR analysis of *Cxcl2* gene expression in 48h TLR2-activated (50ng/mL Pam3CysK4) WT (black bars), IL-10R KO (red bar) or IL-10R/CerS2 DKO (Blue bars) peritoneal macrophage supplemented with CholPC (vehicle), Cer16:0 or Cer24:0 for the last 44h of activation.

All experiments are reported as means ± SD from 3 independent experiments, unless noted otherwise. \**P* < 0.05; \*\**P* < 0.01, \*\*\**P* < 0.005 (two-tailed unpaired Student’s *t* test)

**Supplemental Figure 3:**

A) Flow cytometry-based immune cell profiling of the total CD4 T cells (Lin-/TCRB+/CD4+) or IFNg+ or IL-17A+ CD4 T cells from colonic lamina propria from control (IL-10R Heterozygous), IL-10R KO and IL-10R/CerS2 DKO chimeric mice.

B) Flow cytometry-based immune cell profiling of the colonic lamina propria from WT and CerS2 KO chimera mice

C) scRNAseq UMAP analysis of macrophage markers and inflammatory genes from macrophage from IL-10R KO and IL-10R/CerS2 DKO colon LPs

D) Violin plots of scRNAseq Gene expression analysis from macrophage sorted cells from the colon LP of IL-10R KO or IL-10R/CerS2 DKO chimera mice

All experiments are reported as means ± SD from 3 independent experiments, unless noted otherwise. \**P* < 0.05; \*\**P* < 0.01, \*\*\**P* < 0.005 (two-tailed unpaired Student’s *t* test)

**Supplemental Figure 4:**

A) Heatmap of all sphingolipid metabolism pathway genes from naïve or TLR2 activated WT and IL10 KO BMDMs from RNAseq analysis from samples matched to lipidomics analysis in Figure 1 (TPMs must be above 0 to be listed). Values not scaled.

B) Total ceramides species measured by direct infusion MS from 48h TLR2-activated IL-10 KO BMDMs plus BSA or BSA-18:1 for the last 44h

C) Total, saturated and unsaturated ceramide species measured by direct infusion MS from naïve or 48h TLR2-activated WT BMDMs plus or minus DMSO (vehicle), 10nM SCDi (Cay10566) +/-BSA or 18:1 for the last 44h.

D) qPCR analysis of *Cxcl1* gene expression in 48h TLR2-activated (50ng/mL Pam3CysK4) WT BMDMs plus or minus DMSO (vehicle), 10nM SCDi (Cay10566) +/-BSA or 18:1 for the last 44h.

E) Flow cytometry based immune cell profiling of the colonic lamina propria from Control (LysM Cre+/-) and SCD2 cKO (LysM Cre+/-Scd2^flox/flox^) naïve male mice

F) Flow cytometry based myeloid cell profiling of the colonic lamina propria from Control (LysM Cre+/-) and SCD2 cKO (LysM Cre+/-Scd2^flox/flox^) female mice harvested on day 12 of DSS challenge.

G) Flow cytometry based lymphoid cell profiling of the colonic lamina propria from Control (LysM Cre+/-) and SCD2 cKO (LysM Cre+/-Scd2^flox/flox^) female mice harvested on day 12 of DSS challenge.

All experiments are reported as means ± SD from 3 independent experiments, unless noted otherwise. \**P* < 0.05; \*\**P* < 0.01, \*\*\**P* < 0.005 (two-tailed unpaired Student’s *t* test)

**Supplemental Figure 5:**

A) ELISA analysis of lipocalin from feces of Day 0: heterozygous (het) vs IL-10R KO mice, and Day 14: heterozygous (het) mice gavaged with BSA, IL-10R KO mice gavaged with BSA or IL-10R KO mice gavaged with BSA-24:1 (all male mice)

B) ELISA analysis of lipocalin from feces of Day 0: IL-10 KO female mice versus Day 14: IL-10 KO mice gavaged with BSA or BSA-18:1 (all female mice)

C) Flow cytometry based myeloid cell profiling of the colonic lamina propria from heterozygous (het) mice gavaged with BSA, IL-10R KO mice gavaged with BSA or IL-10R KO mice gavaged with BSA-24:1 (all male mice)

D) Flow cytometry based myeloid cell profiling of the colonic lamina propria from WT mice gavaged with BSA, IL-10 KO mice gavaged with BSA or IL-10 KO mice gavaged BSA-18:1 (all female mice)

All experiments are reported as means ± SD from 3 independent experiments, unless noted otherwise. \**P* < 0.05; \*\**P* < 0.01, \*\*\**P* < 0.005 (two-tailed unpaired Student’s *t* test)

## Notes

### Competing Interest Statement

The authors have declared no competing interest.

