## Supplemental Figures for "IL-10 constrains sphingolipid metabolism via fatty acid desaturation to limit inflammation"

Supplemental Figure 1:

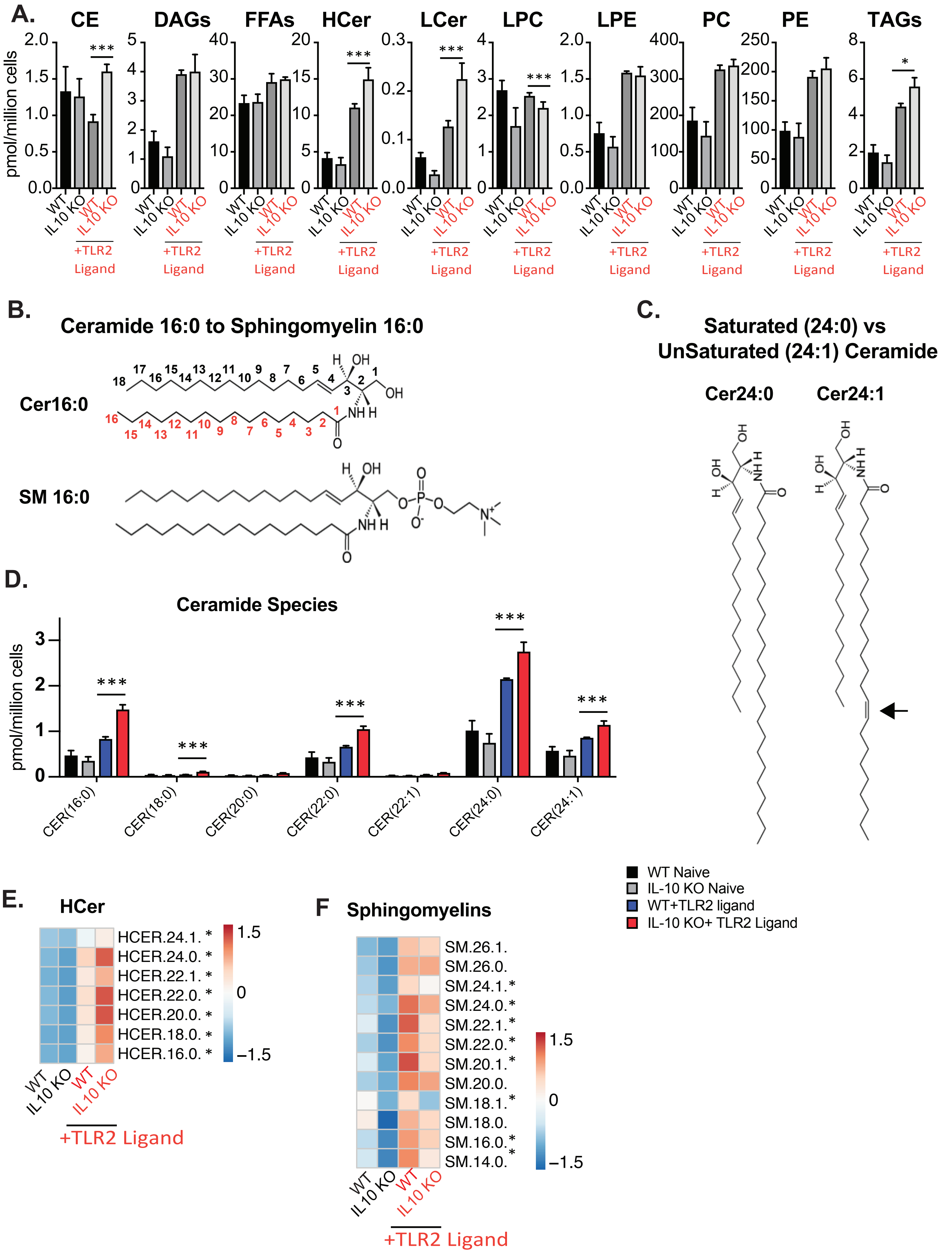

Supplemental Figure 2:

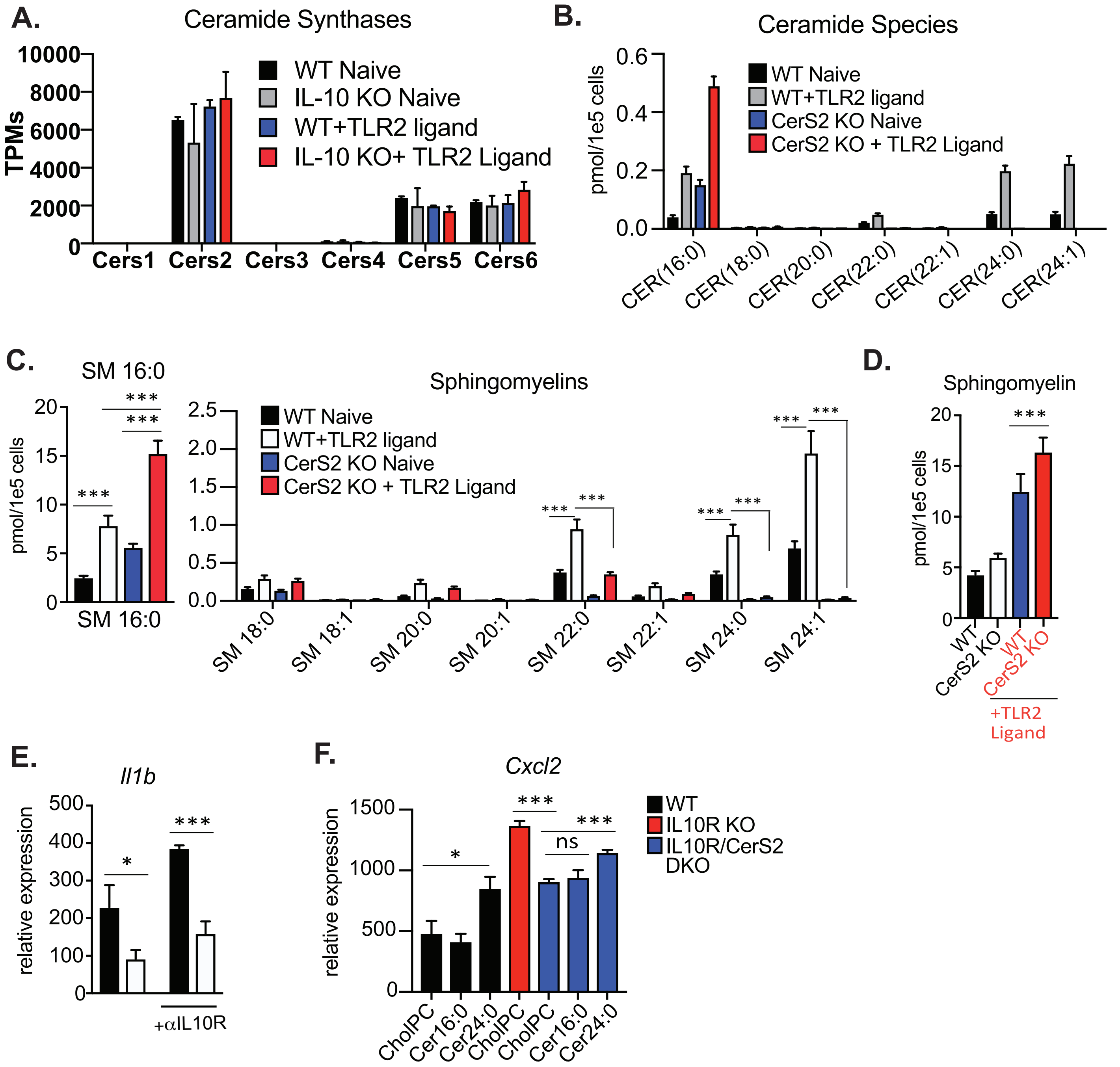

Supplemental Figure 3:

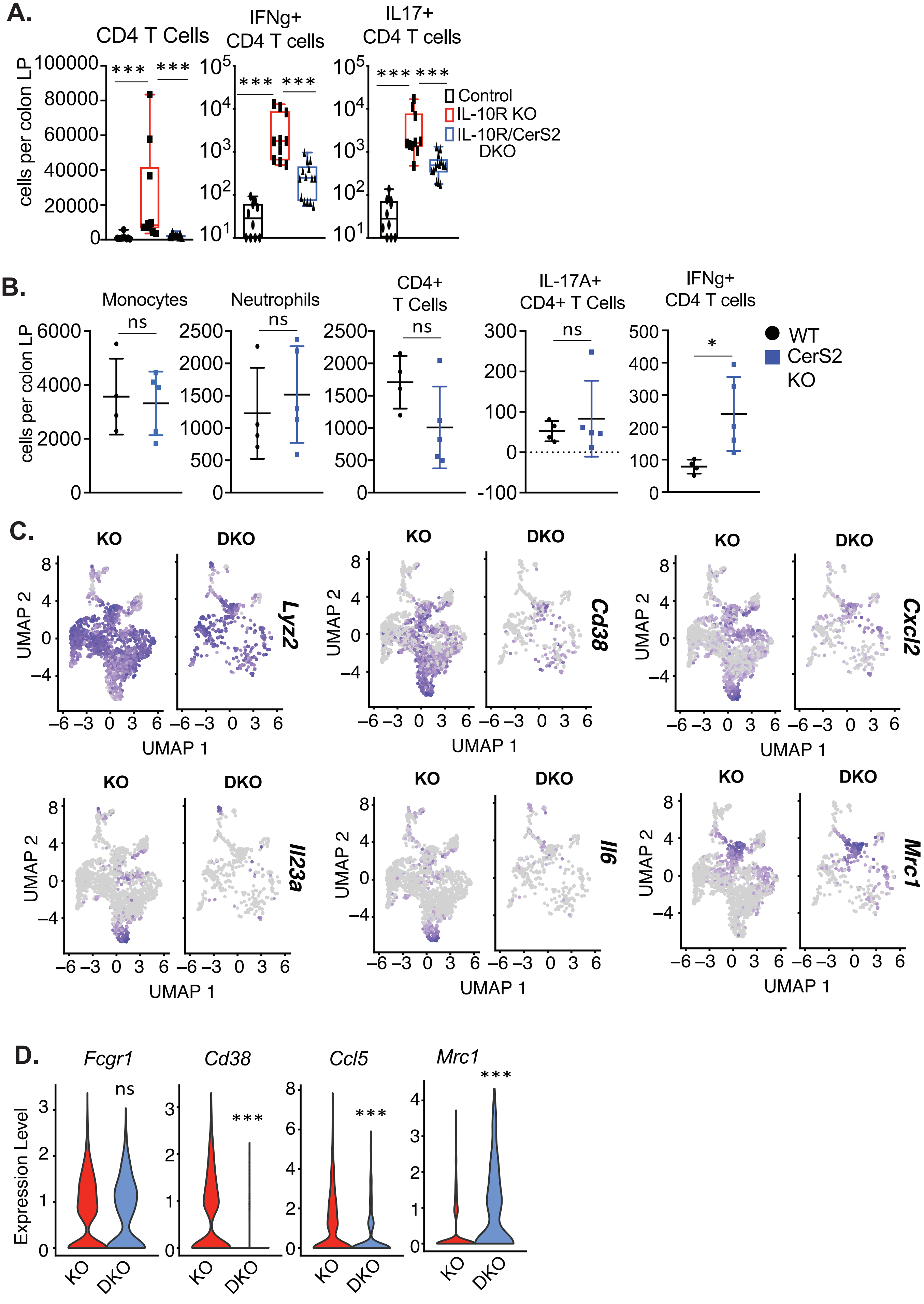

Supplemental Figure 4:

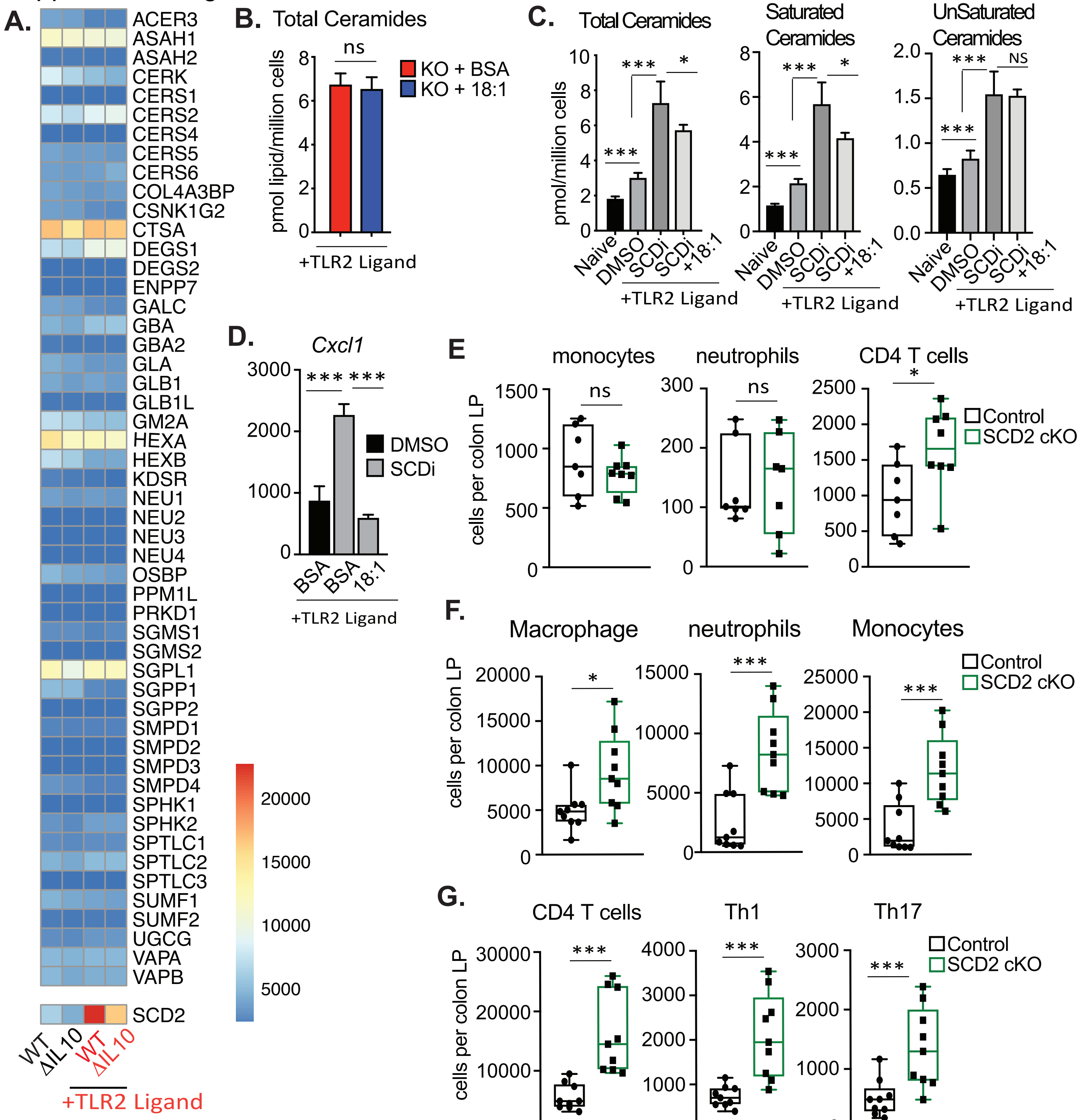

Supplemental Figure 5:

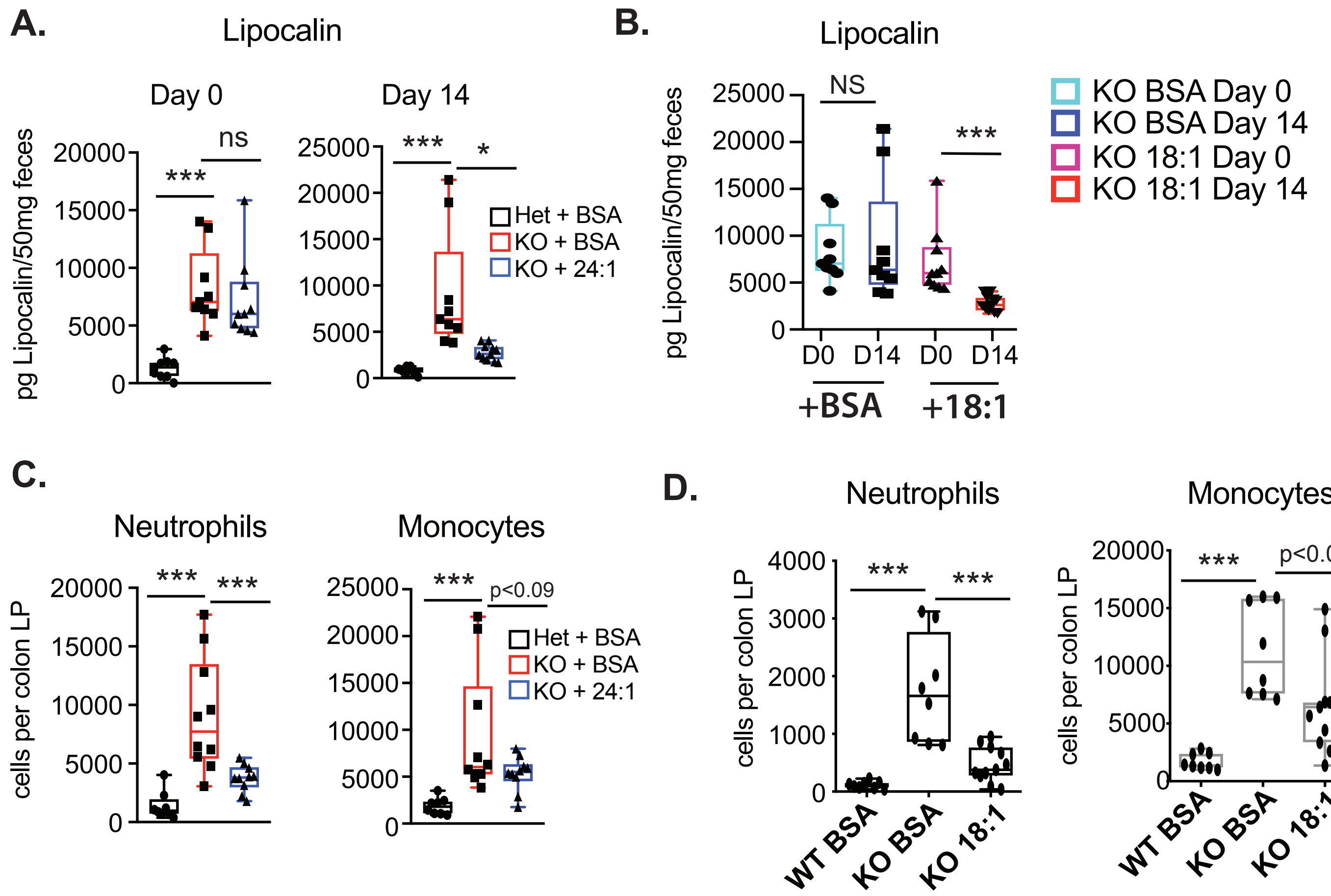
